# Identification of novel genes including *rpmF* and *yjjQ* critical for Type II persister formation in *Escherichia coli*

**DOI:** 10.1101/310961

**Authors:** Shuang Liu, Nan Wu, Shanshan Zhang, Yumeng Zhang, Wenhong Zhang, Ying Zhang

## Abstract

Persister cells, which are characterized by inactive metabolism and tolerance to antibiotics or stresses, pose a significant challenge to the treatment of many persistent infections. Although multiple genes have been reported to be involved in persister formation through transposon mutant library screens, how persisters are formed during the natural process of persister formation as the culture transitions from log phase to stationary phase is unclear. Here, using *E. coli* as a model, we performed a comprehensive transcriptome analysis of gene expression profiles of successive cultures of an *E. coli* culture at different critical time points, starting from persister-free S1-nonexistence phase (3h) to persister appearing S2-emergence phase (4h), and persister abundant stage S3-abundance phase (5h). The differentially expressed genes (≥2-fold) in persister appearing stage (S1 to S2 transition) and persister abundant stage (S1 to S3) were compared, and 51 and 29 genes were identified to be up-regulated, respectively. Importantly, 13 genes (*gnsA, gnsB, ybfA, yjjQ, ymdF, yhdU, csgD, yncN, rpmF, ydcX, yohJ, ssrA, rbsD*) overlap in both persister S2-emergence phase and S3-abundance phase, including a member of the trans-translation pathway (*ssrA*) as well as an orphan toxin (*ydcX*), which are two well-known persister genes while the remaining 11 novel genes (*gnsA, gnsB, ybfA, yjjQ, ymdF, yhdU, csgD, yncN, rpmF, yohJ, rbsD*) have not been reported previously. Persister levels of 7 constructed knockout mutants (*ΔgnsA, ΔybfA, ΔyjjQ, ΔyhdU, ΔcsgD, ΔyohJ* and *ΔrpmF*) and 10 overexpression strains (*gnsA, gnsB, ybfA, yjjQ, ymdF, yhdU, csgD, rpmF, yohJ, rbsD*) in *E. coli* uropathogenic strain UTI89 were determined upon treatment with different cidal antibiotics (ampicillin, levofloxacin and gentamicin). Additionally, ranking of these overlapping genes according to their impact on persister levels were also performed. Two genes (*rpmF* encoding 50S ribosomal subunit protein L32, and *yjjQ* encoding a putative LuxR-type transcription factor) showed the most obvious phenotype on persister levels in both knockout and overexpression studies, which suggests they are broad and key factors for persister formation. While previous studies cannot distinguish if a given persister gene is involved in persister formation or persister survival, our findings clearly identify novel persister forming genes and pathways involving a ribosome protein and a LuxR type transcription factor during the bona fide persister formation process and may have implications for developing improved treatment of persistent infections.

## Introduction

A small percentage of bacterial cells can survive antibiotic or other stress-induced cell death by entering a transiet dormant or slow-growing state (Kint et al., 2012; Lewis, 2010; Zhang, 2014), and this phenomenon is termed persistence and was first discovered by Hobby (1942) and Bigger (1944). Persistence is considered to be associated with chronic and recalcitrant bacterial infections which can ratchet up risks for antibiotic resistance and pose a grave threat to human health (Mulcahy et al., 2010; Zhang et al., 2012; Zhang, 2014; Van den Bergh and Michiels, 2016). Persister cells are believed to form either randomly or as an induced product (Balaban et al., 2004; Nierman et al., 2015; Van den Bergh et al., 2017). Persister cells can be divided into two types, type I persisters and type II persisters. Type I are generated from stationary phase while type II persisters are continuously generated through the whole bacterial growth stage as the culture grows from log phase to stationray phase (Balaban et al., 2004). Multiple molecular mechanisms have been proposed to be involved in the formation of persister (Li and Zhang, 2007; Dorr et al., 2009; Ma et al., 2010; Wang and Wood, 2011; Vega et al., 2012a; Li et al., 2013; Maisonneuve et al., 2013; Marques et al., 2014; Wu et al., 2015) (stringent response, toxin–antitoxin (TA) systems, SOS response, global regulators, signaling molecules, energy metabolism).

Although various persister pathways have been identified, they are mainly studied at one time point of bacterial growth stage, and little is known about the overall dynamic changes of gene expression profile associated with type II persister cell formation naturally during growth cycle from “nonexistence” to “emergence” and finally “abundance”, which we define them as S1, S2, S3, respectively, in this study. Here, using transcriptional analysis by RNA-seq, we provide a “dynamic evolution” of genes when type II persister cells are formed from non-persister cells. We identified many genes that have not been previously reported that are involved in type II persister formation. Fifty-three and thirty-two genes were found to have significantly different expression in S2 and S3 compared with S1, respectively. Specially, 13 genes overlap in both comparison groups. Apart from 4 genes (*gnsB, ymdF, yncN, ybfA*) with uncharacteristic functions and 2 genes (*ssrA, ydcX*) with definite effect on persistence (Li et al., 2013; Islam et al., 2015), the remaining 7 overlapping genes were mapped to ribosomal protein (*rpmF*), regulator of phosphatidylethanolamine synthesis (*gnsA*), membrane protein (*yohJ, yhdU*), DNA-binding transcriptional activator (*csgD*), transcription factor (*yjjQ*) and cytoplasmic sugar-binding protein (*rbsD*). We confirm two genes (*rpmF* and *yjjQ*) play crucial roles in persister formation under multiple antibiotic treatment and stress conditions. Together, our data provide a dynamic profile of genes involved in type II persister formation which leads to novel insight on bona fide mechanisms of persister formation.

## Methods

### Bacteria and culture conditions

*E. coli* K12 strain W3110 from glycerol stocks at −80 °C were 1:1000 diluted in Luria-Bertani (LB) broth (10 g Bacto-tryptone, 5 g yeast extract, and 10 g NaCl/liter) without antibiotics and incubated at 37 °C for 16 hours. For transcriptomics, the overnight culture was diluted 1:10^5^ inoculated in 3.5 liters, 2 liters and 1 liter fresh LB medium for each time point (3 hrs, 4 hrs and 5 hrs) to satisfy the quantity requirement for library construction and grown at 37 °C (100 rpm).

### RNA isolation and preparation of cDNA libraries and RNA-sequencing

Samples of each time point were collected from three independent replicates. Briefly, cells grown to the indicated time points were harvested by centrifugation at 4°C (4000g x 10 min) in 50ml centrifuge tubes and cell cultures were kept on ice during the whole harvest process. Samples were then preserved at −80 °C until RNA was extracted. Total RNA was extracted from the three samples using TRIzol^®^ reagent according to the manufacturer’s instructions (Invitrogen, USA). Integrity of RNA was determined by 2100 Bioanalyzer (Agilent Technologies, USA). RNA was purified using oligo (dT) beads (Illumina, San Diego, CA, USA). Fragmentation buffer was then added to cut the long mRNA into small pieces. The mRNA short fragments were primed with random hexamer primers for synthesis of single-stranded cDNAs which were adopted as the templets for synthesis of the second cDNA strand. The double-stranded cDNAs were purified using the QiaQuick PCR extraction kit (Qiagen). The purified cDNAs were then processed in end repair and appended with poly(A). The appropriate cDNA fragments were obtained through agarose gel electrophoresis and amplified using PCR. The cDNA libraries were sequenced using an Illumina HiSeq™ 2500 (OE Biotech Company, Shanghai, China) and produced 125 bp double-end reads.

### Transcriptome bioinformatic analysis

The raw reads (FastQ format) were qualified by NGS QC Toolkit to abandon end joints, low-quality bases and N-base which resulted in the “clean reads”. All the subsequent analysis was based on the “clean reads”. Reads were mapped to the *E. coli str. K12 substr. W3110* genome (NC_007779.1; NCBI) using Tophat (http://tophat.cbcb.umd.edu/). Then FPKM and count values of the matched reads were obtained using eXpress (Mortazavi et al., 2008). These values were normalized by DESeq (2012) R package (Siska and Kechris, 2017) (estimate Size Factors). Subsequently, P values and fold changes were calculated using nbinomTest. Differential expression genes (DEGs) with P value less than 0.05 were selected for KEGG and COG analysis to determine the gene functions and pathways. Finally, expression profile of DEGs in different samples was displayed using heat map.

### Validation of RNA-seq data by quantitative real-time PCR

Thirty DEGs were chosen randomly and processed in qRT-PCR to validate the RNA-seq data. cDNAs in qRT-PCR were synthesized from the same RNA extractions used for the RNA-seq using a Prime Script RT Reagent Kit (TaKaRa, Japan) and then 1:5 diluted. *rrsB* was selected as the control gene. qRT-PCRs were carried out in ABI 7500 Real-Time instrument thermocycler (Applied Biosystems, Foster City, CA, USA). Relative fold changes were worked out using the 2^−ΔΔCt^ method.

### Knockout mutant construction and persister assay

Deletion of persister genes made in the UTI89 background was achieved by using the λ Red recombination systems, as previously described by Datsenko and Wanner (Datsenko and Wanner, 2000). The deleted genes were stably replaced with a chloramphenicol resistance cassette. All mutants were confirmed with PCR and sequencing (Biosune, Shanghai, China). Persistence was measured by determining the bacterial survival as colony-forming units (CFUs) per 1mL after exposure to cidal antibiotics, i.e., 200 μg/mL ampicillin, or 5μg/mL levofloxacin, or 40 μg/mL gentamicin. Following overnight growth, 1ml cultures were transferred to a 1.5 ml Eppendorf tube and immediately treated with the above antibiotics and incubated at 37°C without shaking for different times. To confirm the persister levels of samples for transcriptome assay, 1 ml culture was withdrawn from the culture of each time point and treated with ampicillin (100 μg/ml) for 3 hours. The cell viability was measured by samples withdrawn at the desired time points, washed and serially diluted in PBS, followed by inoculation onto LB agar without antibiotics. The CFU numbers were counted after overnight incubation at 37°C.

### Construction of overexpression mutant strains

Eleven overlapping genes were amplified by PCR and the primers flanked by *Nco*I and *Eco*RI restriction enzyme sites are listed in Table S1. Following double-enzyme digestion, the resultant DNA fragments were cloned into the same restriction sites of plasmid vector pBAD202, resulting in a series of recombinant plasmids with kanamycin resistance cassette. After selection on kanamycin-containing agar plates and confirmation by DNA sequencing, the resulting constructs were transformed into uropathogenic *E. coli* strain UTI89 and designated as the overexpression mutant strains.

### Measurement of cellular ATP concentrations

The cellular ATP concentrations of wild type and knockout mutants (or overexpression strains) were determined using an ATP assay kit according to the manufacturer’s protocol.

## Results

### Determination of persister levels at three time points under ampicillin treatment

To obtain the gene expression profile of associated with persister formation, the emergence of persisters under ampicillin treatment at different time points was performed. To minimize the influence of type I persister cells in the inoculum, we inoculated the overnight culture in fresh LB broth at a high dilution of 1:10^5^ and incubated in 37°C (100 rpm) for different times. To detect the persister levels, 1 ml bacterial culture from different ages was withdrawn and treated with ampicillin (100 μg/ml) for the same length of time (3 hours). We found no viable cells or persisters existedif the age of cultures was 3hrs or younger. However, when the bacterial culture age was extended to 4 hrs, persister cells started to emerge, but with a number of less than 10 CFU/ml. As we prolonged the incubation time to 5 hrs, there was a surge in the number of persister cells (see Fig. S1). Based on the above results, we defined the three ampicillin-associated critical time points (3hrs, 4hrs and 5hrs) as S1 “nonexistence” to S2 “emergence” and S3 “abundance”, respectively. Bacteria of S1/S2/S3 were then analyzed using RNA-seq to identify the genes differentially expressed in persister emergence and abundance stages S2 and S3 compared with no persister stage S1.

### Identification and verification of differentially expressed genes (DEGs) during persister formation

The up- or down-regulated genes (S2 vs. S1, and S3 vs. S1) whose *p* values were below 0.05 were identified (Table 2). Thirteen overlapping DEGs of S2/S1 and S3/S1 were chosen to validate the reliability of RNA-seq using quantitative real-time PCR (RT-PCR). As a result, all the genes we tested presented concordant expression patterns in RT-PCR and RNA-seq. The results presented here indicate the data from RNA-seq is credible in the follow-up analysis (see Table 1).

**Table 1.** Persister formation related gene transcripts with more than 2-fold up-regulation in S2 compared with S1 (*p*-value <0.05)*.

| NO. | Gene ID | Gene name | Fold change | Gene description |
| --- | --- | --- | --- | --- |
| 1 | 12933889 | *ycdA* | 45.44144 | Uncharacterized protein |
| 2 | **12933782** | ***yjjQ*** | **22.85771** | **Putative transcription factor** |
| 3 | **12933901** | ***ycdX*** | **10.62014** | **Orphan toxin OrtT** |
| 4 | **12930553** | ***ymdF*** | **7.767768** | **Uncharacterized protein** |
| 5 | **12933458** | ***yhdU*** | **7.602703** | **Predicted membrane protein** |
| 6 | **12930933** | ***ybfA*** | **6.672799** | **Uncharacterized protein** |
| 7 | 12932386 | *idnK* | 6.291892 | Pentose phosphate pathway<br>Carbon metabolism |
| 8 | **12933843** | ***gnsA*** | **6.216046** | **Predicted regulator of**<br>**phosphatidylethanolamine**<br>**synthesis** |
| 9 | **12931244** | ***gnsB*** | **5.887559** | **Protein GnsB** |
| 10 | 12934174 | *yfcV* | 5.534535 | Uncharacterized fimbrial-like<br>protein |
| 11 | **12930501** | ***yncN*** | **4.272272** | Hypothetical protein |
| 12 | **12933181** | ***yohJ*** | **4.326215** | **Conserved inner membrane**<br>**protein** |
| 13 | 12933927 | *zntA* | 3.713078 | Lead, cadmium, zinc and<br>mercury-transporting ATPase |
| 14 | 12930672 | *idnD* | 3.620335 | L-idonate 5-dehydrogenase<br>(NAD(P)(+)) |
| 15 | 12932329 | *arsR* | 3.140484 | Arsenical resistance operon<br>repressor \|sequence-specific<br>DNA binding transcription<br>factor activity |
| 16 | 12933817 | *ykgE* | 3.131381 | Predicted oxidoreductase |
| 17 | 12930745 | *yadK* | 3.087688 | Uncharacterized fimbrial-like<br>protein |
| 18 | 12934032 | *yebN* | 3.006995 | Conserved inner membrane<br>protein |
| 19 | 12930936 | *ybfH* | 2.8713 | Hypothetical protein |
| 20 | 12930776 | *fadE* | 2.87019 | Fatty acid degradation\|Fatty acid<br>metabolism |
| 21 | 12931540 | *yfcY* | 2.823285 | Valine, leucine and isoleucine<br>degradation\|Fatty acid<br>degradation |
| 22 | 12932494 | *torR* | 2.789674 | Two-component system |
| 23 | 12934145 | *fsaB* | 2.738138 | Fructose-6-phosphate aldolase 2 |
| 24 | 12934116 | *yhhI* | 2.71125 | Predicted transposase |
| 25 | 12934023 | *insC* | 2.667615 | IS2 insertion element repressor<br>InsA |
| **26** | **12931070** | ***csgD*** | **2.621622** | **DNA-binding transcriptional**<br>**activator** |
| 27 | 12934464 | *rbsA* | 2.598083 | ABC transporters |
| 28 | 12931901 | *yfdY* | 2.588437 | Predicted inner membrane<br>protein |
| 29 | 12932146 | *arsC* | 2.586667 | Arsenate reductase; |
| 30 | 12933846 | *torT* | 2.487179 | Periplasmic sensory protein<br>associated with the TorRS<br>two-component regulatory<br>system |
| 31 | 12933202 | *fxsA* | 2.485491 | Inner membrane protein |
| 32 | 12931368 | *yobB* | 2.467409 | Uncharacterized protein YobB; |
| 33 | 12933149 | *glmU* | 2.440769 | Amino sugar and nucleotide<br>sugar metabolism |
| 34 | 12931356 | *torY* | 2.427427 | Cytochrome c-type protein<br>TorY; |
| 35 | 12930508 | *ldhA* | 2.387548 | Pyruvate metabolism |
| **36** | **12932242** | ***rbsD*** | **2.384947** | **ABC transporters** |
| 37 | 12934234 | *pspC* | 2.338203 | Transcriptional activator |
| 38 | 12931784 | *ybeF* | 2.21888 | Uncharacterized HTH-type<br>transcriptional regulator YbeF; |
| **39** | **12932035** | ***rpmF*** | **2.21342** | **50S ribosomal protein L32** |
| 40 | 12932624 | *yfiA* | 2.213267 | Cold shock protein associated<br>with 30S ribosomal subunit |
| 41 | 12931638 | *yjfV* | 2.174975 | Arsenical pump membrane<br>protein |
| 42 | 12931815 | *ibpB* | 2.160241 | Small heat shock protein IbpB; |
| 43 | 12933322 | *ygeP* | 2.115695 | Uncharacterized protein YgeP; |
| 44 | 12932504 | *yagI* | 2.105242 | Uncharacterized HTH-type<br>transcriptional regulator YagI; |
| **45** | **12930217** | ***ssrA*** | **2.103644** | **Trans-translation** |
| 46 | 12931840 | *yohK* | 2.063606 | Inner membrane protein YohK; |
| 47 | 12933596 | *ibpA* | 2.062559 | Small heat shock protein IbpA; |
| 48 | 12933928 | *yhhP* | 2.058133 | Uncharacterized protein |
| 49 | 12931415 | *cobS* | 2.05437 | Porphyrin and chlorophyll<br>metabolism |
| 50 | 12932416 | *ymcB* | 2.026644 | Uncharacterized protein |
| 51 | 12932000 | *degP* | 2.019632 | Two-component system |

\* The bold fonts represent the 13 overlapping genes.

**Table 2.**
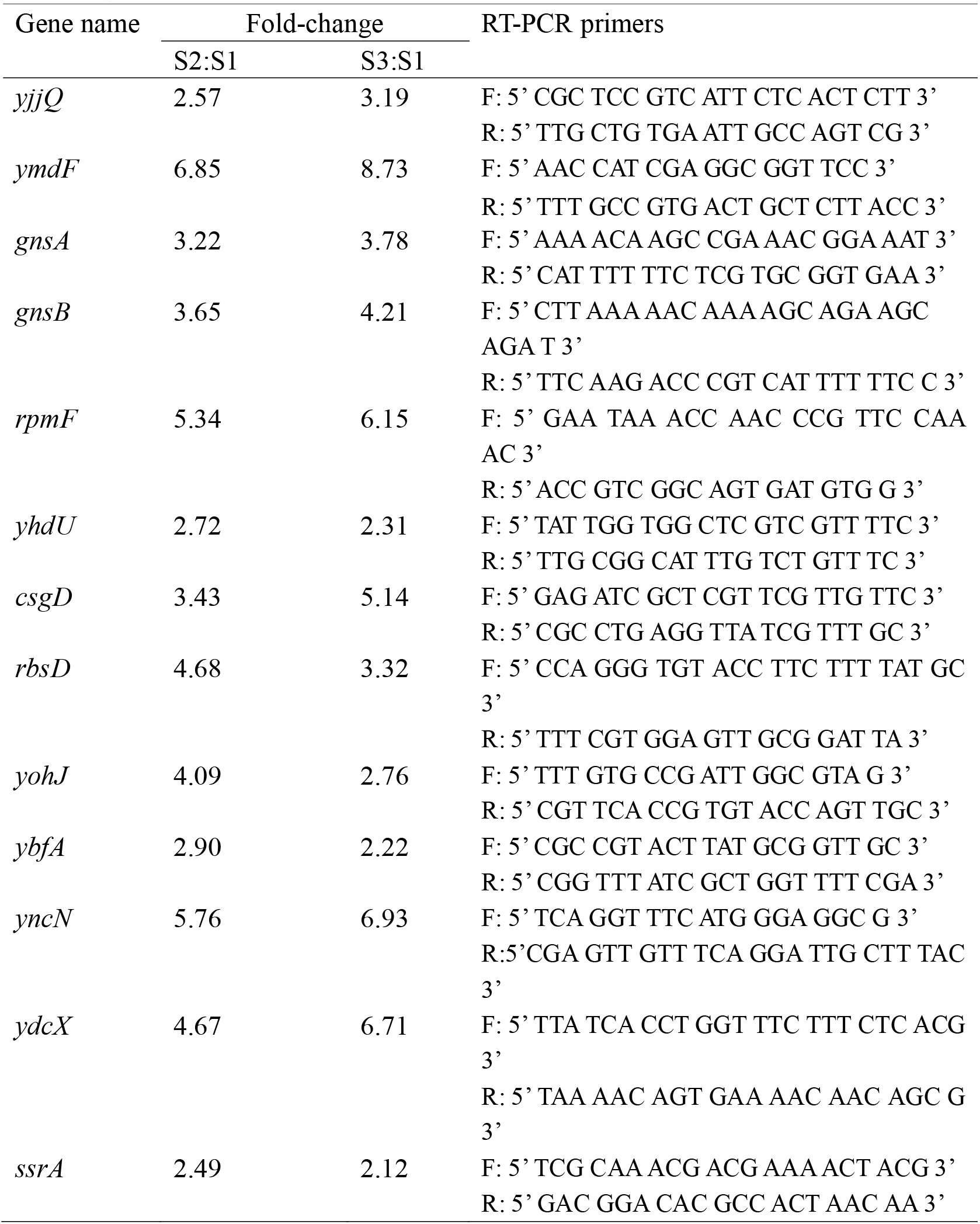
Fold-changes of the expression levels of the 13 genes that overlap in S2:S1 and S3:S1 by RT-PCR

### Core DEGs in S2 versus S1 and S3 versus S1

S1 “nonexistence” to S2 “emergence” and S3 “abundance” referred to the three different stages of persister formation, therefore DEGs of S2 vs. S1 and S3 vs. S1 would allow us to discover critical genes in the formation of persisters. Fifty-one and twenty-nine DEGs that showed significantly elevated levels (≥2-fold increase) were identified based on the values of S2/S1 and S3/S1 comparison, respectively (see Table 2). Nine of the 51 genes up-regulated in S2 vs. S1 encoded uncharacterized or hypothetical proteins (*ydcA, ymdF, ybfA, gnsB, yncN, ybfH, yobB, ygeP, yhhP, ymcB*). Of the remaining genes, 7 were assigned to metabolism pathways according to the KEGG analysis and they are phosphatidylethanolamine synthesis (gnsA), fatty acid metabolism (fadE, *yfcY*), carbon metabolism (*idnK*), porphyrin and chlorophyll metabolism (*cobS*), amino sugar and nucleotide sugar metabolism (*glmU*) and pyruvate metabolism (*ldhA*), respectively. Other pathways such as trans-translation (*ssrA*), ATP-binding cassette (ABC) transporters (*rbsA, rbsD*), and two-component system (*torR, torT, degP*) were also involved in the formation of persisters. In addition, genes encoding toxin (*ydcX*), ribosomal protein or ribosomal associated proteins (*rpmF, yfiA*), phage shock protein (*pspC*), small heat shock proteins (*ibpB, ibpA*), membrane proteins (*yhdU, yohJ, yebN, yfdY, fxsA, yohK, yfjV*), transcriptional activators or factors (*yjjQ, csgD, arsR, pspC, ybeF, yagI*), many enzymes: ATPase (*zntA*), dehydrogenase (*idnD*), arsenate reductase (*arsC*), oxidoreductase (*ykgE*), transposase (*yhhI*) and fructose-6-phosphate aldolase 2 (*fsaB*). Moreover, fimbrial-like protein (*yfcV, yadK*), IS2 insertion element repressor (*insC*), cytochrome c-type protein (*torY*) also showed significant increase in S2 compared with S1 (see Fig. 1A). Of these genes, phage shock protein (*pspC*) has been reported to be involved in indole-mediated persister formation (Vega et al., 2012b). And *ydcX* also participates in persister formation as an orphan toxin (Islam et al., 2015), and *ssrA* involved in trans-translation has previously been shown to be involved in persister formation (Li et al., 2013).

**Figure 1.**
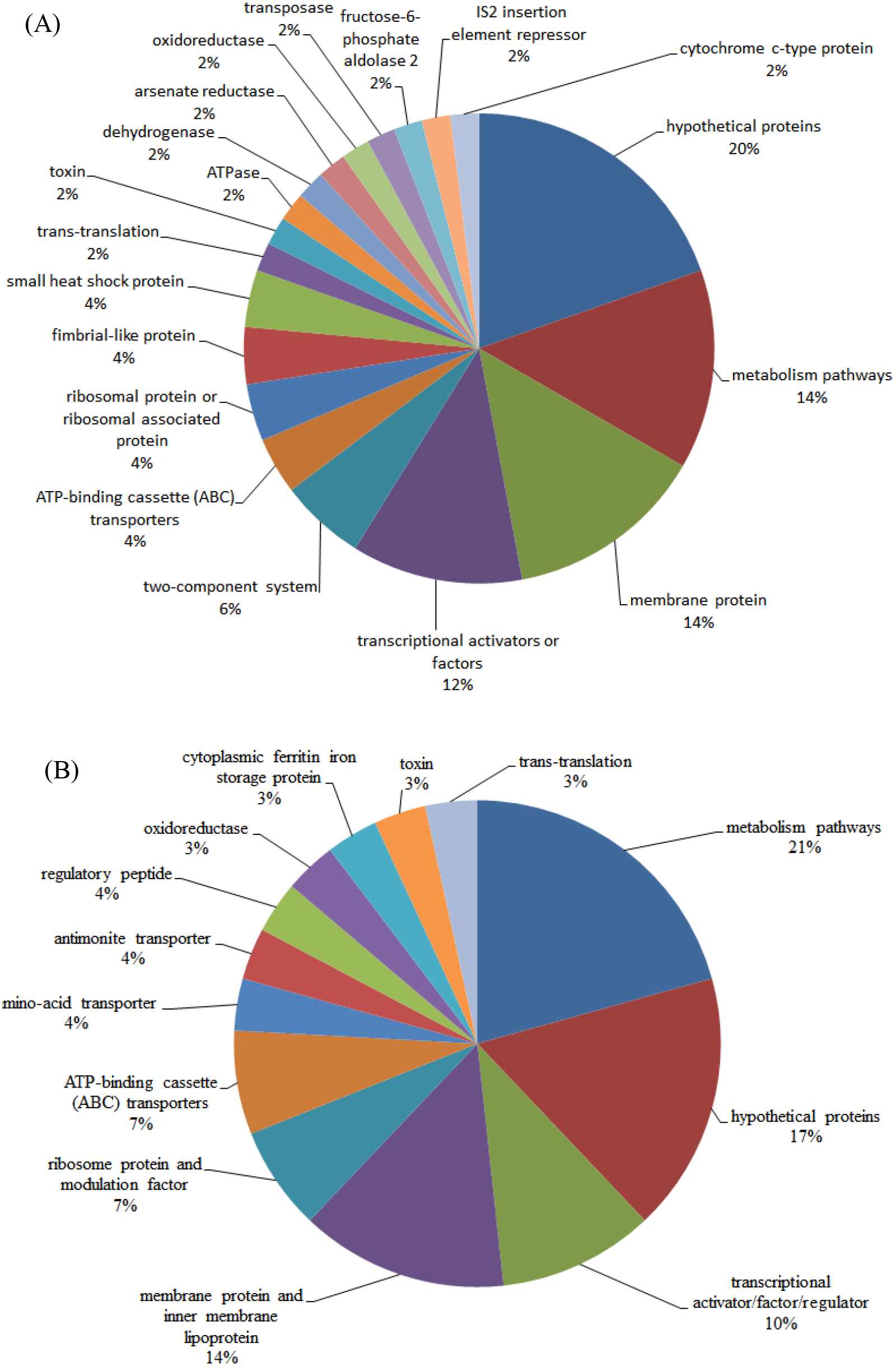
Persister associated DEG (≥2-fold increase) distributions and percentage. (A) S2:S1 and (B) S3:S1

During the period of “nonexistence to abundance”, the number of DEGs (≥2-fold increase) was only 29 (see Table 3). Thirteen of them overlapped with those of S2/S1 while the other 16 genes encode proteins that are involved in arginine and proline metabolism (*argC, argG, argD*), glycerophospholipid metabolism (*glpD*), propanoate metabolism (*tdcD*), DNA-binding transcriptional regulator (*envR*), ABC transporters (*ccmD*), amino-acid transporter (*yeeF*), antimonite transporter (*arsB*), ribosome modulation factor (*rmf*), regulatory peptide (*mokB*), oxidoreductase (*ymjC*), membrane protein (*yohM*), inner membrane lipoprotein (*yghJ*), cytoplasmic ferritin iron storage protein (*ftn*) and uncharacterized protein (*yhfG*) (see Fig. 1B). As for the overlapping genes, 4 genes (*ymdF, ybfA, yncN, gnsB*) encode putative or uncharacterized proteins. Other overlapping genes were related to ribosomal protein (*rpmF*), toxin (*ydcX*), membrane proteins (*yohJ, yhdU*), DNA-binding transcriptional activator or transcriptional factors (*csgD, yjjQ*), trans-translation (*ssrA*), phosphatidylethanolamine synthesis (*gnsA*) and ABC transporters (*rbsD*). The 13 overlapping genes of S2/S1 and S3/S1 were supposed to be of significance for both “boot-up” and “shoot-up” of persister formation and required a thorough investigation. Apart from two genes (*ssrA* and *ydcX*) which have been known to participate in the persister formation (Li et al., 2013; Islam et al., 2015), the other 11 overlapping genes have not been reported to be related to persistence.

**Table 3.** Persister formation related gene transcripts with more than 2-fold up-regulation in S3 compared with S1 (*p*-value <0.05) *.

| NO. | Gene ID | Gene name | Fold change | Gene description |
| --- | --- | --- | --- | --- |
| **1** | **12930501** | ***yncN*** | **15.67** | **Hypothetical protein** |
| 2 | 12931025 | *rmf* | 15.6 | Ribosome modulation factor |
| **3** | **12932035** | ***rpmF*** | **15.2** | **50S ribosomal protein L32** |
| 4 | 12933891 | *mokB* | 14.6 | Regulatory peptide |
| 5 | 12932090 | *yhfG* | 14.4 | Uncharacterized protein YhfG |
| **6** | **12930553** | ***ymdF*** | **10.1** | **Uncharacterized protein** |
| **7** | **12933901** | ***ydcX*** | **6.6** | **Orphan toxin OrtT** |
| **8** | **12933458** | ***yhdU*** | **5.5** | **Predicted membrane protein** |
| **9** | **12933782** | ***yjjQ*** | **5.0** | **LuxR-type transcription factor** |
| **10** | **12930933** | ***ybfA*** | **4.2** | **Uncharacterized protein** |
| 11 | 12932958 | *envR* | 4.0 | DNA-binding transcriptional regulator |
| **12** | **12930217** | ***ssrA*** | **3.8** | **Trans-translation** |
| 13 | 12932271 | *glpD* | 3.5 | Glycerophospholipid metabolism |
| 14 | 12931490 | *ccmD* | 3.2 | ABC transporters |
| **15** | **12933181** | ***yohJ*** | **3.06** | **Conserved inner membrane protein** |
| 16 | 12933196 | *argC* | 3.0 | Arginine and proline metabolism\|2-Oxocarboxylic acid metabolism\|Biosynthesis of amino acids |
| **17** | **12932242** | ***rbsD*** | **2.9** | **ABC transporters** |
| 18 | 12933881 | *ymjC* | 2.6 | Predicted oxidoreductase |
| 19 | 12931423 | *yeeF* | 2.6 | Predicted amino-acid transporter |
| 20 | 12933985 | *yghJ* | 2.5 | Predicted inner membrane lipoprotein |
| **21** | **12931244** | ***gnsB*** | **2.5** | **Protein GnsB** |
| 22 | 12934127 | *argG* | 2.5 | Alanine, aspartate and glutamate metabolism\|Arginine and proline metabolism\|Biosynthesis of amino acids |
| 23 | 12933423 | *tdcD* | 2.4 | Propanoate metabolism |
| 24 | 12931961 | *yohM* | 2.1 | Membrane protein conferring nickel and cobalt resistance |
| 25 | 12934013 | *ftn* | 2.1 | Cytoplasmic ferritin iron storage protein |
| 26 | 12931748 | *argD* | 2.0 | Lysine biosynthesis\|Arginine and proline metabolism\|2-Oxocarboxylic acid metabolism\|Biosynthesis of amino acids |
| **27** | **12931070** | ***csgD*** | **2.0** | **DNA-binding transcriptional activator** |
| **28** | **12933843** | ***gnsA*** | **2.0** | **Predicted regulator of phosphatidylethanolamine synthesis** |
| 29 | 12932331 | *arsB* | 2.0 | Arsenite/antimonite transporter |

\* The bold fonts represent the 13 overlapping genes.

### Persister levels of the overlapping gene knockout mutants in the presence of ampicillin, gentamicin and levofloxacin

Because *ssrA* and *ydcX* have previously been shown to be involved in persister formation (Li et al., 2013; Islam et al. 2015), we focused on assessing the impact of the knockout strains on persister levels of the remaining 11 of the 13 previously unreported overlapping genes under ampicillin treatment. However, only 7 knockout mutants (*ΔgnsA, ΔybfA, ΔyjjQ, ΔyhdU, ΔcsgD, ΔyohJ* and *ΔrpmF*) were successfully constructed presumably because some genes are either essential, or lack of sequence homolog (i.e. *yncN*) in UTI89 strain. We chose to construct the mutants in uropathogenic strain UTI89 because it is ureopathogenic strain that is conducive to further in-vivo experiments. Cells of wild type and knockout mutants from log phase (~10^8^ CFU/ml) were challenged with ampicillin (200 μg/ml), gentamicin (40 μg/ml) and levofloxacin (5 μg/ml). Persister numbers were determined at 3 hrs, 5 hrs and 24 hrs, respectively. Of the 7 knockout mutants tested, *rpmF*, which encodes the 50S ribosomal subunit protein L32, manifested the most obvious phenotype in persister assays with all three antibiotics. After 24 hrs of ampicillin exposure, the *ΔrpmF* mutant had ~10^5^-fold decrease from the initial CFU numbers (10^8^ CFU/ml) while UTI89 had 10^4^-fold decrease from 10^8^ CFU/ml (see Fig. 2A). When exposed to gentamicin and levofloxacin, *ΔrpmF* also exhibited lower persister levels compared with UTI89 (10~100-fold change compared with UTI89) (see Fig. 2B and C). *ΔybfA*, also had dramatic decreased persister levels (10~1000-fold decrease) in the presence of ampicillin as well as gentamicin compared with UTI89. While under levofloxacin treatment, *ΔybfA* showed a negligible effect on persister levels. *ΔgnsA*, though exhibited significantly lower persister levels compared with UTI89 (10~10^3^-fold decrease) when exposed to gentamicin and levofloxacin, showed a higher persister level (>10-fold increase) in the presence of ampicillin. *ΔyhdU* also showed a 100-fold increase in persister numbers when exposed to ampicillin. However, under gentamicin and levofloxacin treatment, *ΔyhdU* exerted little impact on persister numbers. Other knockout mutant strains, including *ΔcsgD* and *ΔyohJ*, exhibited similar persister levels to UTI89 under ampicillin and levofloxacin treatment. However, persister levels of *ΔcsgD* and *ΔyohJ* were dramatically decreased when exposed to gentamicin (~10^6^-fold decrease for *ΔcsgD* and ~10-fold decrease for *ΔyohJ*). However, when tested in the presence of levofloxacin and gentamicin, instead of higher persister level, *ΔyhdU* showed the similar persister number as UTI89 and *ΔgnsA* exhibited 10~100-fold reduction in persister level. We did not observe any alteration of persister numbers for *ΔyjjQ* compared with the wild type under our experiment conditions. The impact of *ybfA* exerted on persister formation was specific to ampicillin and gentamicin while *ΔcsgD* and *ΔyohJ* affected persister formation only in the presence of gentamicin. Contrary to our expectation, our data revealed two unique knockout mutants (*ΔgnsA* and *ΔyhdU*) which displayed 10~100-fold higher persister numbers than UTI89 when exposed to ampicillin. And *ΔgnsA* even exerted opposite effect on persister formation under different antibiotic treatment. This seemingly paradoxical result observed for *ΔgnsA* was not unexpected, instead, it may indicate that its role is to suppress persister formation. Overall, the data presented above revealed only one knockout mutant, *ΔrpmF* mutant exhibited a universal effect on persister formation for all the antibiotics tested.

**Figure 2.**
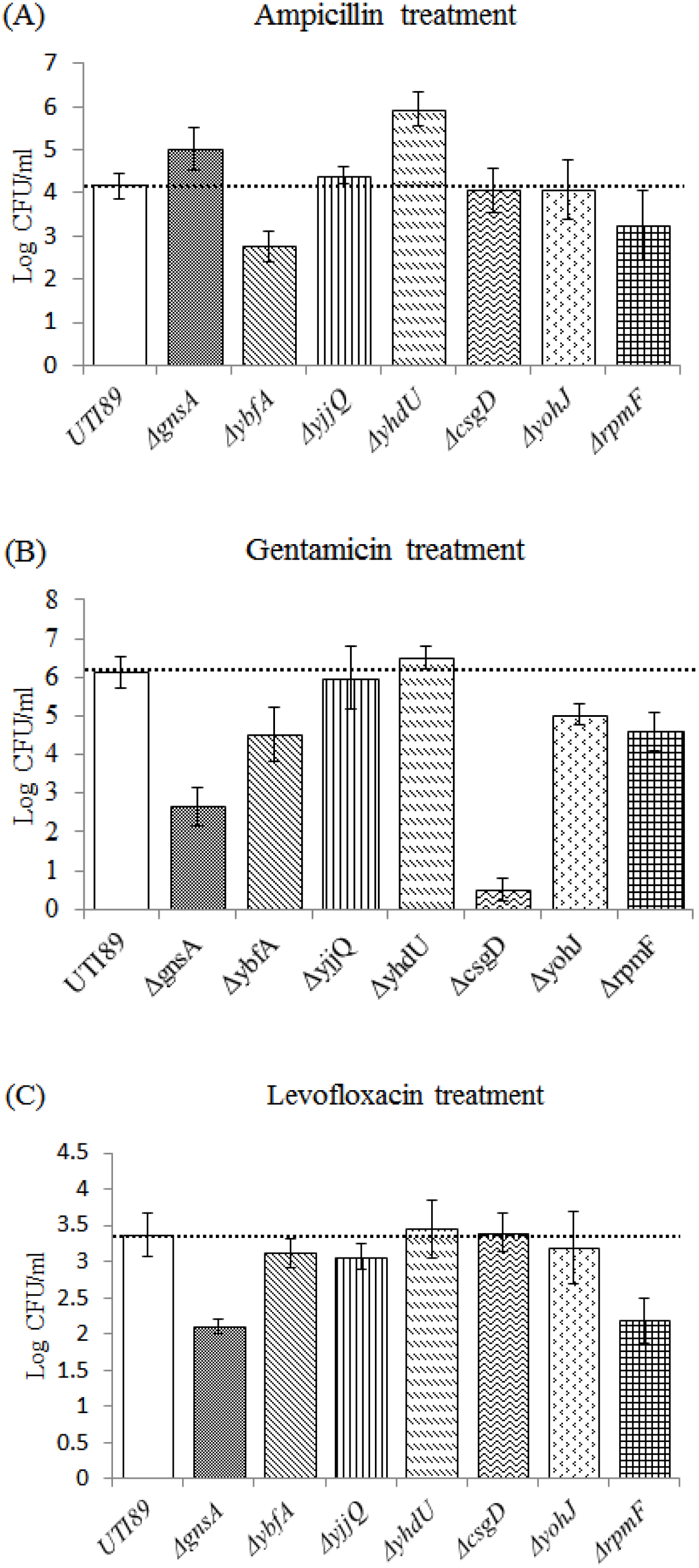
Effect of the 7 overlapping gene knockout mutations on *E. coli* persister formation. Cultures of UTI89 and persister gene knockout mutants grown to log phase and were immediately treated with (A) ampicillin (200 μg/ml) for 24 hours, (B) gentamicin (40 μg/ml) for 5 hours, (C) levofloxacin (5 μg/ml) for 3 hours. Surviving bacteria were counted after 16h incubation at 37 °C on LB plates without antibiotics.

### Effect of overexpression of the 10 overlapping genes on persister levels

To further investigate whether these genes participate in persister formation, we constructed 10 overexpression strains and subjected them to treatment with ampicillin (200 μg/ml), levofloxacin (5 μg/ml) and gentamicin (50 μg/ml). UTI89 transformed with pBAD202 plasmid was used as a control. pBAD202 plasmids containing the 10 genes were constructed and 0.2% arabinose was added to induce the gene expression. Twenty-four hours after ampicillin treatment, 7 overexpression strains (*gnsA, gnsB, yhdU, csgD, yohJ, rpmF* and *rbsD*) had 10~10^4^ -fold higher persister levels than the control strain (see Fig. 3A). Contrary to the RNA-seq data, two genes (*yjjQ* and *ymdF*) exhibited negative effect on persister formation (see Fig. 3A), while *ybfA* showed no influence on persister levels (see Fig. 3A). Collectively, the overexpression study under ampicillin treatment further confirmed the validity of the transcriptome assay, though two genes (*yjjQ* and *ymdF*) showed the opposite effect. These overexpression strains were also exposed to gentamicin and levofloxacin. Fig. 3B shows in the presence of gentamicin, 7 overexpression mutants (*gnsB, yhdU, csgD, rpmF, rbsD, yjjQ* and *ymdF*) displayed consistent effect with what they did when exposed to ampicillin. Overexpression of the 5 genes could lead to 10~100-fold increase in persister formation while *yjjQ* and *ymdF* still had negative effect on persister formation (100~1000-fold decrease compared with UTI89+pBAD202). *gnsA*, though could increase persister level by more than 100-fold, had negative effect on persister formation (~2.5-fold decrease) when exposed to gentamicin. Overexpression of *ybfA* which had little effect on persister level under ampicillin treatment could increase persister numbers by 4-fold in gentamicin treatment. *yohJ* had little influence on persister formation. When exposed to levofloxacin, increased persister levels compared with the control strain were observed for only 3 overexpression strains (*gnsA, ybfA* and *ymdF*), with ~4-fold for *ybfA* and >10-fold for *gnsA* and *ymdF* (see Fig. 3C). Other genes, except for *yjjQ* and *yohJ* which exhibited >100-fold and ~4-fold decreased persister levels, had no obvious effect on persister formation. Notably, in the presence of levofloxacin, only two genes (*gnsA* and *yjjQ*) had the same effect on persister formation as observed in the presence of ampicillin. Collectively, our data suggest that of the genes *gnsA, gnsB, ybfA*, *yhdU, csgD, yohJ, rpmF* and *rbsD* conductive to persister formation, 5 genes (*gnsB, yhdU, csgD, rpmF* and *rbsD*) were specific to ampicillin and gentamicin, one gene (*gnsA*) was ampicillin- and levofloxacin-specific, one gene (*ybfA*) was gentamicin- and levofloxacin-specific and one gene (*yohJ*) was specific only to ampicillin. Of the genes (*yjjQ*, *ymdF* and *yohJ*) which exhibited negative role in persister formation, *yjjQ* had a general effect for all the three antibiotics, while *ymdF* was specific to ampicillin and gentamicin, and *yohJ* was specific to levofloxacin.

**Figure 3.**
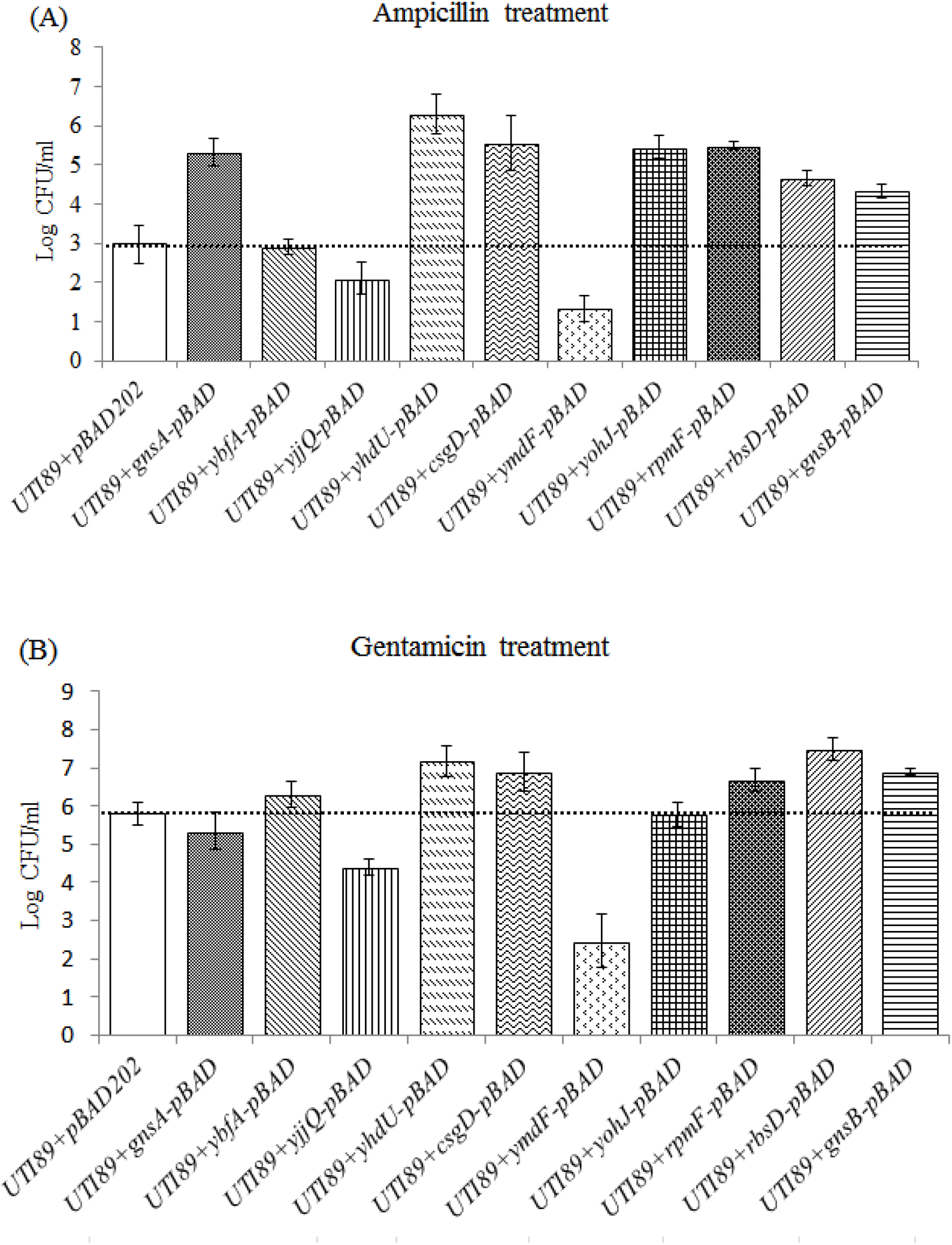
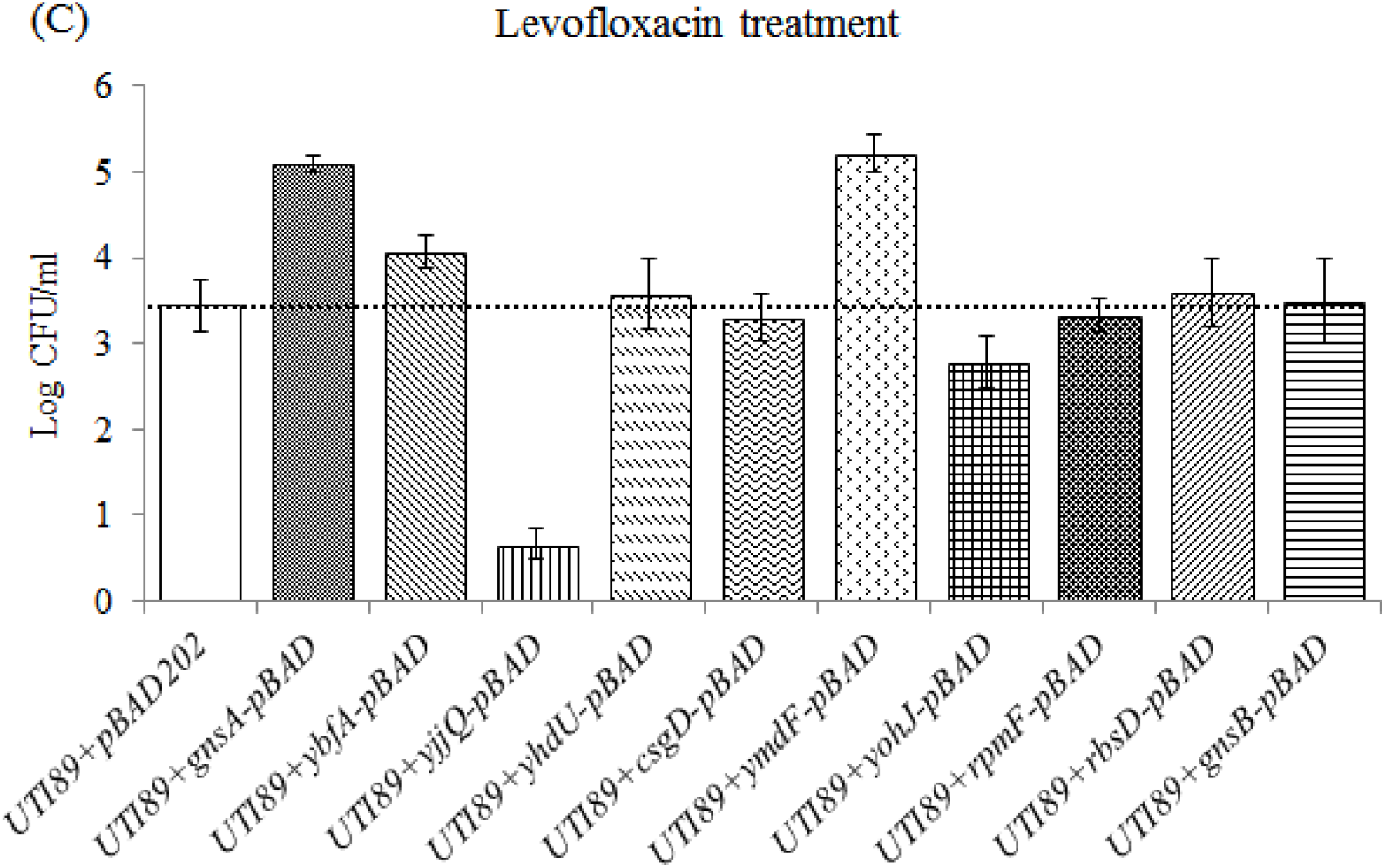
Effect of overexpression of the 10 overlapping genes on *E. coli* persister formation. Cultures of UTI89+pBAD202 and persister gene overexpression mutants grown to log phase and were immediately treated with (A) ampicillin (200 μg/ml) for 24 hours, (B) gentamicin (40 μg/ml) for 5 hours, (C) levofloxacin (5 μg/ml) for 3 hours. Surviving bacteria were counted after 16h incubation at 37 °C on LB plates without antibiotics.

### Ranking of the overlapping genes according to their impact on persister levels under antibiotic exposure

To determine the relative importance of the 10 overlapping genes (*gnsA, gnsB, ybfA, yjjQ, ymdF, yhdU, csgD, rpmF, yohJ, rbsD*) under rxposure to all antibiotics, the results (≥5-fold change) from knockout and overexpression assay were gathered and ranked, respectively. Among the 8 knockout mutants, *rpmF* was shown to play a key and broad role in the process of persister formation with 3 points under treatment with all the three antibiotics. The second most important genes were *ybfA* and *gnsA* with 2 scores. Although *gnsA* exhibited significant effect on persister formation under all the three antibiotics, its knockout mutant showed an enhancement in persister level under ampicillin treatment, which has an opposite effect in the presence of gentamicin and levofloxacin. *yohJ, csgD* and *yhdU* scored only 1 point, which suggests that their impact on persister formation is limited to only one antibiotic. Unfortunately, *yjjQ* scored 0 in its knockout mutant (see Table 4). Table 5 shows a distinct gene arrangement for the 10 overexpression strains. *yjjQ* overexpression strain showed a broad impact on persister formation and scored 3 points. Overexpression strains of seven genes (*ymdF, gnsA, gnsB, rpmF, yhdU, csgD* and *rbsD*) made 2 points. Among these genes, *ymdF* showed significant effects on persister levels upon treatment with all the three antibiotics. However, the effect *ymdF* exerted in the presence of ampicillin and gentamicin was opposite to that in the presence of levofloxacin. *yohJ* received 1 point only when exposed to ampicillin while overexpressing *ybfA* and *yncN* resulted in similar persister levels to the control strain. Furthermore, we also observed some gene knockout mutants exhibited a different persistence profile when compared with their overexpression strains.

**Table 4.** Ranking of the top 7 persister genes according to their knockout mutant scores upon exposure to antibiotics^a^.

| Knockout genes | Pathways or proteins | Scores for each antibiotic <sup>b</sup> |  |  |  | Sum |
| --- | --- | --- | --- | --- | --- | --- |
|  |  | Amp | Gen | Lev | Opposite |  |
| *rpmF* | 50S ribosomal protein L32 | 1 | 1 | 1 |  | 3 |
| *ybfA* | Hypothetical protein | 1 | 1 | 0 |  | 2 |
| *gnsA* | predicted regulator of phosphatidylethanolamine synthesis | 1 | 1 | 1 | -1 | 2 |
| *yohJ* | Conserved inner membrane protein | 0 | 1 | 0 |  | 1 |
| *csgD* | DNA-binding transcriptional activator in two-component regulatory system | 0 | 1 | 0 |  | 1 |
| *yhdU* | Predicted membrane protein | 1 | 0 | 0 |  | 1 |
| *yjjQ* | Putative transcription factor | 0 | 0 | 0 |  | 0 |

<sup>a</sup> Genes were ranked according to cell survival under exposure to three different antibiotics. The genes whose overexpression strains show difference ( $\geq 5$ -fold) compared to the parent strain are scored as “1” point, whereas those mutants that show difference ( $< 5$ -fold) were scored as “0”. All scores for a given mutant are calculated from the sum of different antibiotic exposures to obtain the ranking order of the persister genes.
<sup>b</sup> The three abbreviations “Amp”, “Gen” and “Lev” are symbols for “Ampicillin”, “Gentamicin” and “Levofloxacin”, respectively. “Opposite” represents that a given mutant exhibited decreased persister levels in the presence of one or two antibiotics while showing increased persister numbers in other antibiotic exposures compared with UTI89.

**Table 5.** Ranking of the 10 persister related genes according to their overexpression scores upon exposure to antibiotics^a^.

| Overexpression genes | Pathways or proteins | Scores for each antibiotic <sup>b</sup> |  |  |  | Sum |
| --- | --- | --- | --- | --- | --- | --- |
|  |  | Amp | Gen | Lev | Opposite |  |
| *yjjQ* | Putative transcription factor | 1 | 1 | 1 |  | 3 |
| *ymdF* | Hypothetical protein | 1 | 1 | 1 | -1 | 2 |
| *gnsA* | Predicted regulator of phosphatidylethanolamine synthesis | 1 | 0 | 1 |  | 2 |
| *gnsB* | Hypothetical protein | 1 | 1 | 0 |  | 2 |
| *rpmF* | 50S ribosomal protein L32 | 1 | 1 | 0 |  | 2 |
| *yhdU* | Predicted membrane protein | 1 | 1 | 0 |  | 2 |
| *csgD* | DNA-binding transcriptional activator in two-component regulatory system | 1 | 1 | 0 |  | 2 |
| *rbsD* | Predicted cytoplasmic sugar-binding protein | 1 | 1 | 0 |  | 2 |
| *yohJ* | Conserved inner membrane protein | 1 | 0 | 0 |  | 1 |
| *ybfA* | Hypothetical protein | 0 | 0 | 0 |  | 0 |

<sup>a</sup> Genes are ranked according to cell survival under exposure to three different antibiotics. The genes whose overexpression strains show difference ( $\geq 5$ -fold) compared to the parent strain are scored as “1” point, whereas those mutants that show difference ( $< 5$ -fold) are scored as “0”. All scores for a given mutant are calculated from the sum of different antibiotic exposures to obtain the ranking order of the persister genes.
<sup>b</sup> The abbreviations “Amp”, “Gen” and “Lev” are symbols for “Ampicillin”, “Gentamicin” and “Levofloxacin”, respectively. “Opposite” represents that a given strain exhibited decreased persister levels in the presence of one or two antibiotics while showing increased persister numbers in exposure to other antibiotics compared with UTI89.

### Relative ATP concentration assay

Since ATP levels correlate with persister levels, we also analysed the ATP levels for knockout or overexpression strains. Figure 4A and B show the relative ATP levels for 6 knockout mutants (*Δ csgD, ΔgnsA, Δ yohJ, Δ ybfA, Δ yhdU, Δ rpmF*) and 9 overexpression strains (*yohJ, csgD, yhdU gnsA, gnsB, rbsD, rpmF, yjjQ, ymdF*). Of the 6 knockout mutants, interestingly, ATP production dramatically increased in the *ΔrpmF* (5.9-fold change). ATP produced in *ΔyhdU* was significantly lower than that in UTI89 (the ATP ratio was 0.42). The other 4 knockout mutants (*ΔcsgD, ΔgnsA, ΔybfA, ΔyohJ*) produced similar ATP levels as UTI89. Of the 9 overexpression strains, only *yjjQ* overexpression exhibited a similar ATP concentration as UTI89+pBAD202 vector control strain and all the other 8 overexpression strains (*yohJ, csgD, yhdU gnsA, gnsB, rbsD, rpmF, ymdF*) had lower ATP concentrations than UTI89+pBAD202, indicating their role in persistence.

**Figure 4.**
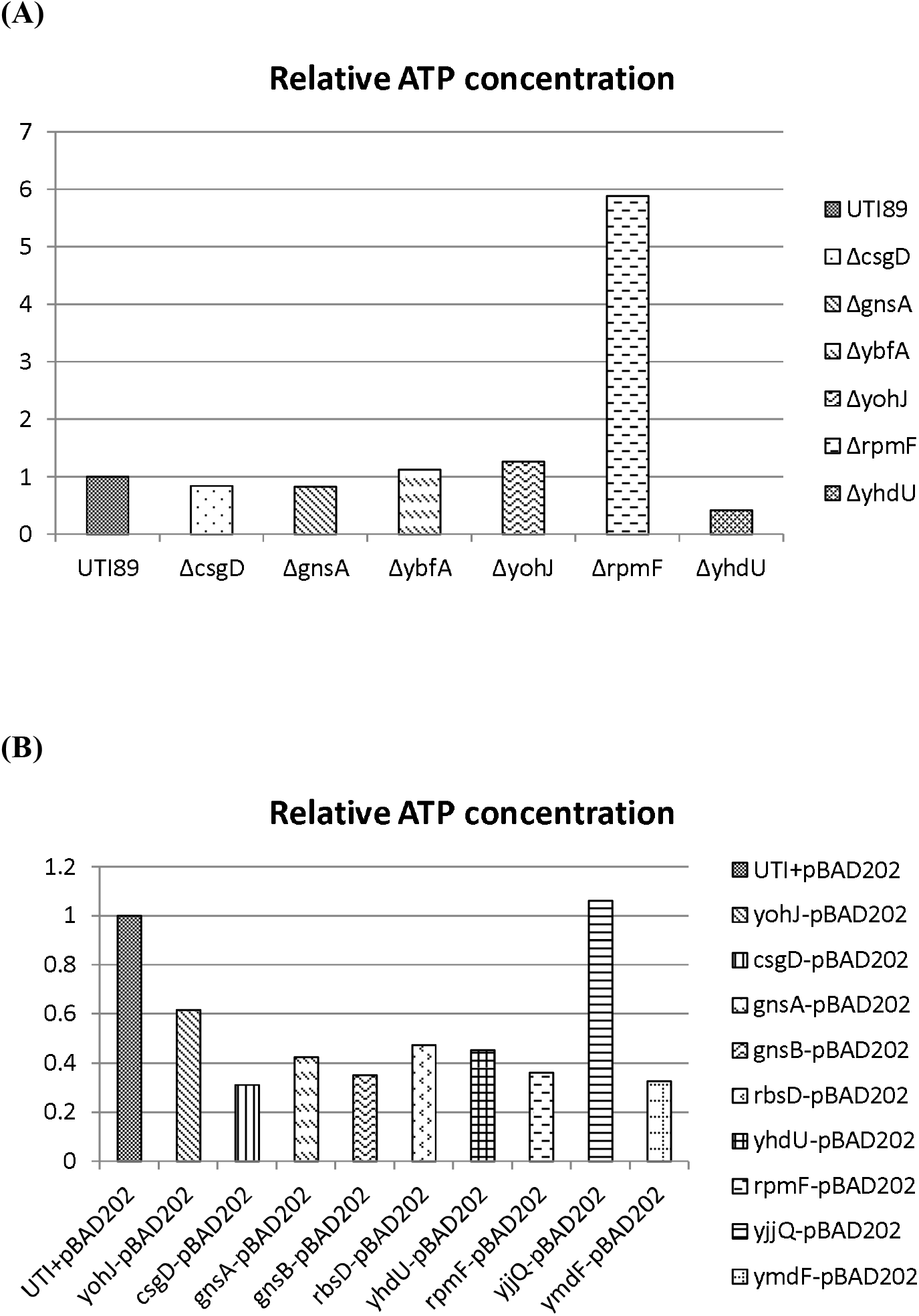
The relative ATP concentrations of 6 knockout mutants and 9 overexpression strains. Cultures of knockout mutants (A) and overexpression strains (B) grown to log phase and immediately subjected to ATP assay as described in Methods.

## Discussion

In this study, we completed the detection of dynamic transcriptional profiles in the process of persister formation for *E. coli* K12 strain W3110 using Illumina RNA sequencing technology. Numerous previous studies investigated mechanisms involved in drug specific persistence by comparison the differentially expressed genes of samples from “pre-drug treatment” and “post-drug treatment”. Samples used here were not treated by antibiotics but had specific antibiotic tolerance profiles. By drawing a comprehensive blueprint of the gene expression levels for three key points (S1 “nonexistence”, S2 “emergence” and S3 “abundance”) of persister formation under ampicillin treatment, we observed 51 and 29 genes were significantly up-regulated (>2-fold change) in S2 and S3 in comparison with S1. We mainly focused on the up-regulated genes and differences between S2 and S1 or S3 and S1 because the period from S1 to S2 (S3) was a “boot-up” (“shoot-up”) process and we believe the two periods (S1 to S2 and S1 to S3) are of critical importance. As stated above, different pathways were involved in persister “boot-up” and “shoot-up” stages. However, metabolic pathways, membrane proteins and transcription factors occupied the principal parts in both stages. This suggests during persister formation, there is a good chance that bacteria undergo considerable changes in their metabolic state as well as membrane characteristics. This concept appears to contradict the current acceptable view that persisters are predominantly dormant and in agreement with Orman and Brynildsen who claim “Dormancy is not necessary or sufficient for bacterial persistence”(Orman and Brynildsen, 2013). This is mainly because we are looking at genes involved in the actual process of persister formation rather than when persisters are already formed. Further experiments should be performed to detect the effects of these metabolic pathways on ATP production. Membrane and membrane proteins have previously been shown to participate in persistence (Cui et al., 2016; Guo et al., 2017), and our data indicate that their roles in persister formation may be far more important and complicated than previously thought, and changes to lowermembrane permeability to drugs has been noted in some dormant bacteria (Dick T, 2015)

By comparing the 51 DEGs in S2/S1 and the 29 DEGs in S3/S1, we observed 13 DEGs (*gnsA, gnsB, ybfA, yjjQ, ymdF, yhdU, csgD, yncN, rpmF, ydcX, yohJ, ssrA, rbsD*) that overlapped between S2/S1 and S3/S1. The 13 overlapping DEGs may play crucial roles throughout the persister formation process. It is surprising that only two previsouly known persister genes *ssrA* (Li and Zhang, 2013) and *ydcX* () are identifed in our study despite many persister genes have been reported previously. This result further confirmed the significance of *ssrA* and *ydcX* in persister formation. However, this did not mean other persister genes were less important than them. A plausible explanation is that other previously identified persister genes either displaytheir effect on persister formation at different conditions rather than the natural persister formation as a function of ageing or time as in this study, or their influence is more on persister maintenance or survival rather than persister formation, as is investigated here.

We found that *rpmF* is involved in participation in multidrug tolerance. *rpmF*, which encodes 50S ribosomal subunit protein L32, is responsible for protein synthesis. Since persisters are considered slow-growing or in dormant state, and metabolically inactive, this prompted us to hypothesize that deletion of *rpmF* would have a significant effect on metabolic proteins, disturb the homeostasis of metabolism in bacteria leading to decreased persister levels. Another explanation is that deletion of *rpmF* would enhance the activity of drug or stress targets and increase the susceptibility of *ΔrpmF*. The multidrug and multiple stress susceptibility of *ΔrpmF* have previously been shown to be hypersensitive to ampicillin, sulfamethoxazole, rifampicin and metronidazole (Tamae et al., 2008). However, subinhibitory concentrations of antibiotics were used in that study, but were not subjected to persister assays where the antibiotic concentrations are usually far more than MICs. Our study is the first to demonstrate the role of *rpmF* in persister formation. Since RpmF is a conserved protein, it is likely that such homologs play similar roles in persister formation in other bacteria. Future studies are needed to confirm this and address the mechanism of RpmF in persister formation. RpmF could serve as a good persister drug target for future drug development.

*rpmF* and *plsX* gene encoding a protein involved in membrane lipid synthesis and several fatty acid biosynthetic genes (*fabH, fabD* and *fabG*) are cotranscribed in *E. coli*. Organization of these genes into an operon may play a role in the coordinated regulation of the synthesis of ribosomes, cell membranes and fatty acid and lipid biosynthesis (Podkovyrov S1, Larson TJ. 1995) and link these processes to cellular metabolism. It is of interest to note that the same fatty acid synthesis operon *fabHDG* is activated by FadR but inhibited by ppGpp, a well-known molecule mediating persister formation (My L, et al. 2013). In addition, it has been shown that *rpmF* is involved in biofilm formation in *Actinobacillus pleuropneumoniae* (Grasteau A, et al., 2011) and that *rpmF* (L32) mutant is hypersensitive to ROS-generating agent hydroxyurea (HU) (Nakayashiki T, Mori H, 2013). These findings are consistent with our findings that RpmF involvement in persister formation may be mediated via ppGpp. Future studies are needed to address the detailed mechanism.

In summary, the data presented here provides a portrait of the overall profile of genes involved in type II persister formation. In particular, besides two known genes including a member of the trans-translation pathway (*ssrA*) and an orphan toxin (*ydcX*), 11 novel genes (*gnsA, gnsB, ybfA, yjjQ, ymdF, yhdU, csgD, yncN, rpmF, yohJ, rbsD*) previously not reported are identified in this study. Among them, two genes *rpmF* (encoding 50S ribosomal subunit protein L32) and *yjjQ* (a LuxR-type transcription factor) are identified as two key factors for persister formation. Our findings shed new ligth on mechanisms of type II persister formation and provide new therapeutic targets for intervention.

---

**Table S1.**
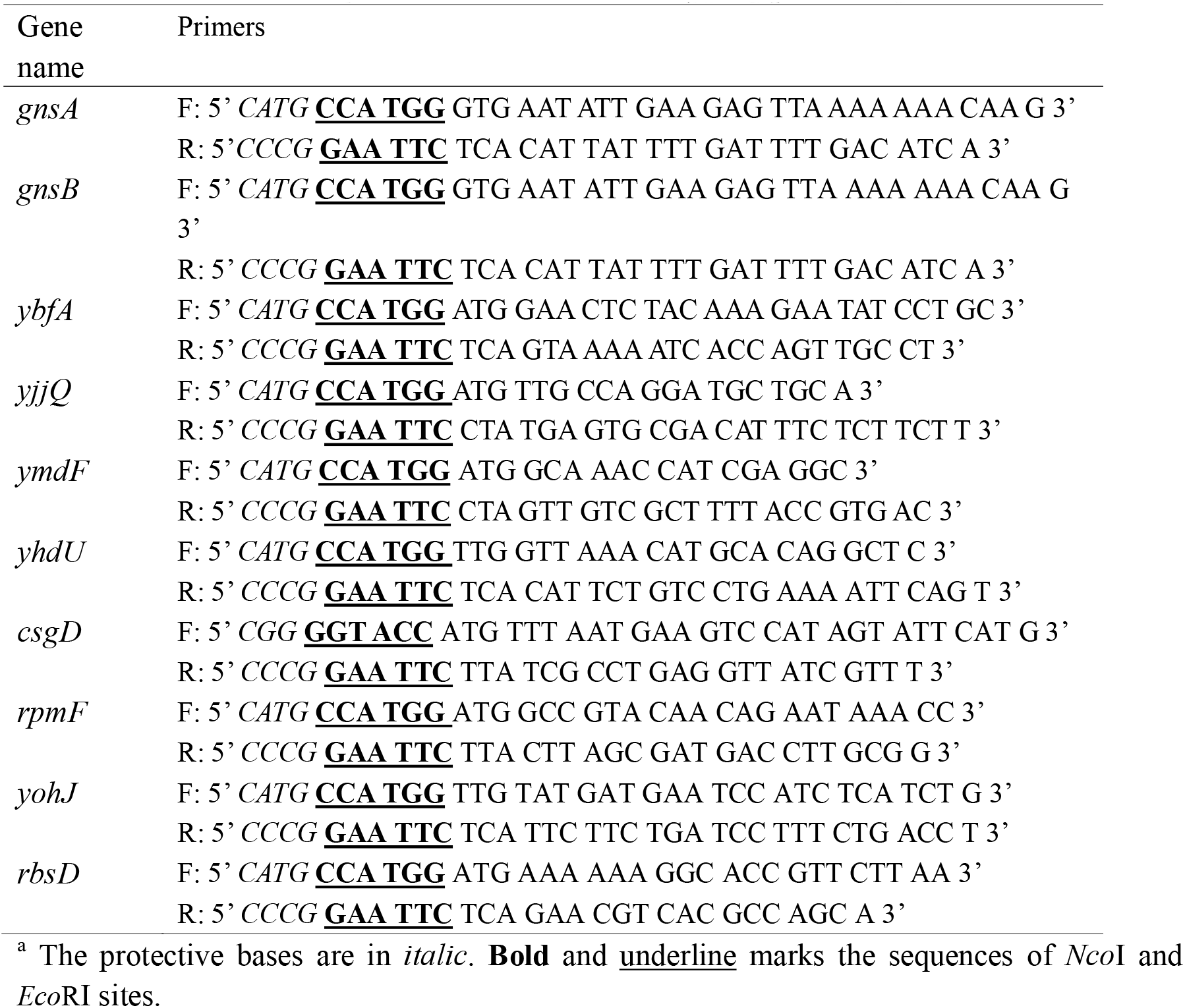
Primers for amplification of the 10 overlapping persister genes^a^

## Supplementary Figure 1

**Figure S1.**
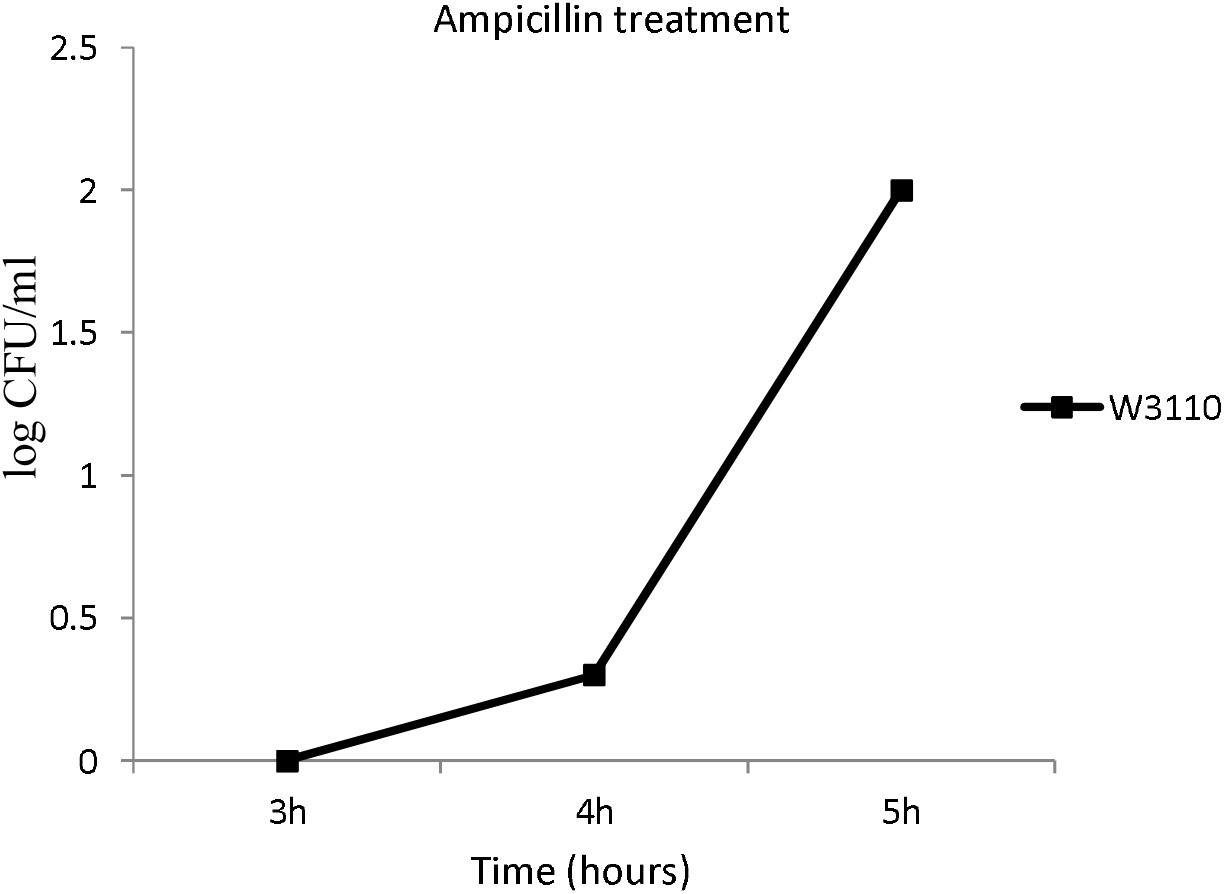
Different persister levels of *E. coli* at three time points. Cultures of *E. coli* W3110 grown to different time points (3h-S1, 4h-S2 and 5h-S3) were withdrawn and immediately treated with ampicillin (100 μg/ml) for 3 hours. Surviving bacteria were counted after 16h incubation at 37 °C on LB plates without antibiotics. As can be seen, no persisters are formed at 3 hrs (S1) but persisters begin to appear at 4 hrs (S2) and shoot up considerably at 5hrs (S3).

